# Assessment of batch-correction methods for scRNA-seq data with a new test metric

**DOI:** 10.1101/200345

**Authors:** Maren Büttner, Zhichao Miao, F Alexander Wolf, Sarah A Teichmann, Fabian J Theis

## Abstract

Single-cell transcriptomics is a versatile tool for exploring heterogeneous cell populations. As with all genomics experiments, batch effects can hamper data integration and interpretation. The success of batch effect correction is often evaluated by visual inspection of dimension-reduced representations such as principal component analysis. This is inherently imprecise due to the high number of genes and non-normal distribution of gene expression. Here, we present a k-nearest neighbour batch effect test (kBET, <u>https://github.com/theislab/kBET</u>) to quantitatively measure batch effects. kBET is easier to interpret, more sensitive and more robust than visual evaluation and other measures of batch effects. We use kBET to assess commonly used batch regression and normalisation approaches, and quantify the extent to which they remove batch effects while preserving biological variability. Our results illustrate that batch correction based on log-transformation or *scran* pooling followed by *ComBat* reduced the batch effect while preserving structure across data sets. Finally we show that kBET can pinpoint successful data integration methods across multiple data sets, in this case from different publications all charting mouse embryonic development. This has important implications for future data integration efforts, which will be central to projects such as the Human Cell Atlas where data for the same tissue may be generated in multiple locations around the world.

[Before final publication, we will upload the R package to Bioconductor]

## Introduction

The term “batch effect” is used to describe variation that emerges through technical effects that arise when samples are handled in distinct batches. Usually, this situation occurs if one repeats an experiment with biologically equivalent cells (e.g. different patients of the same disease) or technically equivalent cells (e.g. sequencing cells of the same culture condition on subsequent days) as depicted in **Fig. 1a**. Both biological and technical variations contribute substantially to the total variability in single-cell RNA-sequencing (scRNA-seq) data. In a balanced design for a sequencing experiment, we are able to identify and distinguish biological from technical variation (**Fig. 1b**). In contrast, a confounded design groups cells of the same condition into the same sequencing runs, and separates biologically distinct cells into entirely distinct handling and sequencing runs. This confounds biological with technical variability. In such a setup, a worst case scenario might be that the technical variation swamps out the biological variation. Furthermore, differences between replicates in scRNA-seq data can arise from different sequencing depths: fewer genes are detected at shallow sequencing depths compared to deeper sequencing^1,2^.

**Fig. 1:**
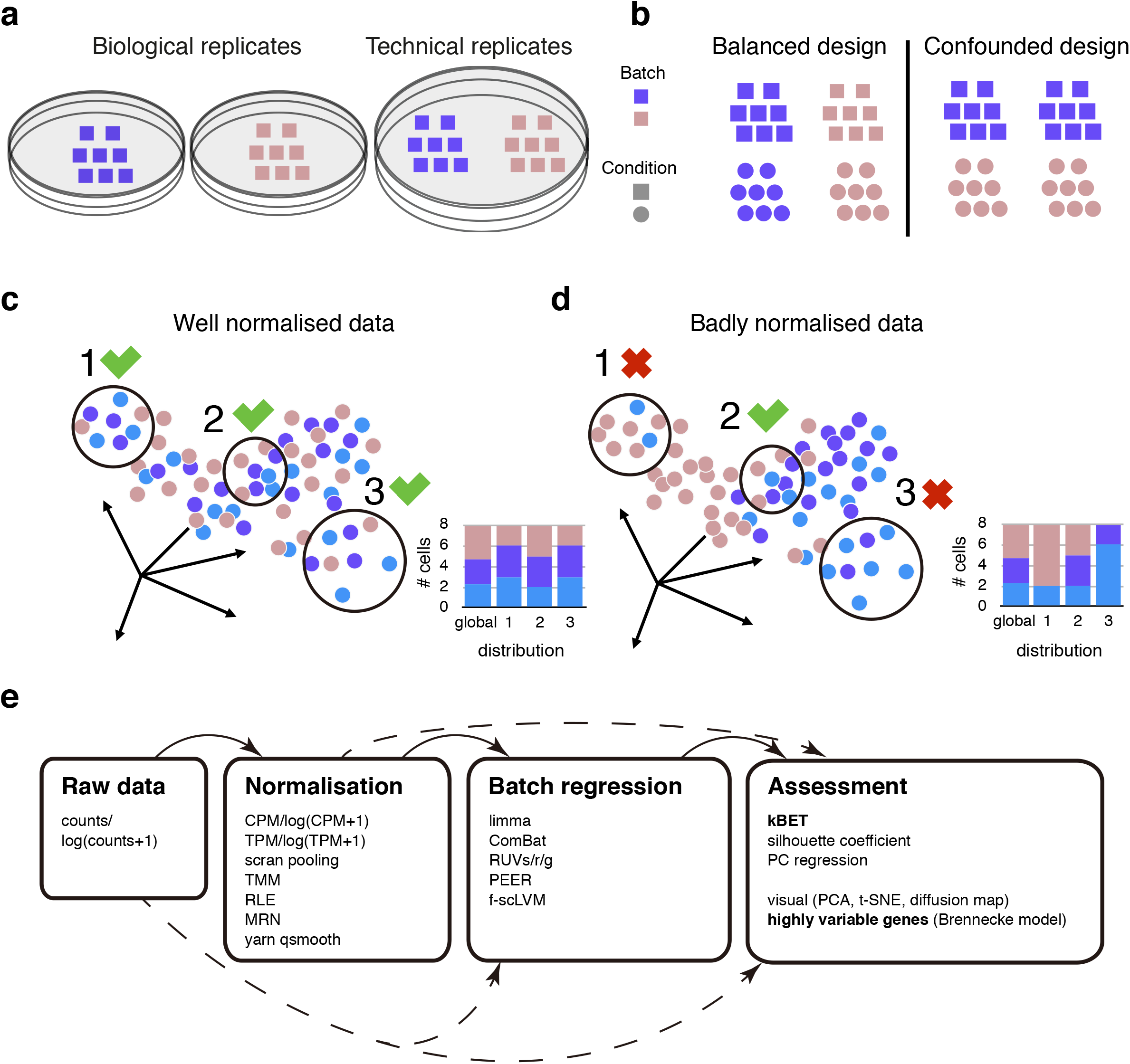
Batch types and the concept of kBET. Estimating the batch effect in single-cell RNA sequencing data. **a)** Biological and technical replicates have different origins. Technical replicates are derived from the same biological samples (in this case cell cultures), while biological replicates are independent samples. **b)** Experimental designs: A balanced design allows one to separate technical and biological sources of variation, while a confounded design mixes both. c) and d) illustrate the concept of kBET. **c)** In a data set with replicates without batch effects, the fraction of the batch label does not differ from the global label distribution in any neighbourhood. **d)** If a data set has a batch effect, data points from respective batches tend to cluster with their ‘peers’. Then the fraction of batch labels differs considerably in arbitrarily chosen neighbourhoods. **e)** Overview of normalisation and batch regression methods as well as assessment approaches.

Several methods have been proposed to remove or reduce batch effects in single-cell data while preserving biological variability. In general, we can distinguish spike-in based and non-spike-in based methods^3,4^. A spike-in is a mix of synthetic, poly-adenylated RNAs as, for example, designed by the External RNA Control Consortium (ERCC) that do not map to the reference genome and that cover different lengths, GC-contents and initial concentrations. However, technical variance of ERCC spike-ins can differ from the variation of biological samples^5^. Hence, the standardisation of data by spike-ins does not necessarily always reflect variation in endogenous mRNA content^6^. Also, droplet-based experimental setups such as DropSeq^7^, inDrop^8^ or the commercial platform Chromium (10x Genomics) do not allow the use of spike-ins. Hence, methods that ‘match’ several replicates without a reference concentration are more practical and preferable. Examples of such methods are factor analysis based methods such as removal of unwanted variation (RUV)^5^ and probabilistic estimation of expression residuals (PEER)^9^. Other non-spike-in approaches use downsampling^10–12^ of the reads or cell-specific scaling factors from pooling across samples^13^ for normalisation. We provide an overview to single-cell normalisation and batch correction methods in **Supplementary Table 1**.

Given the wide variety of normalisation and batch correction strategies available, we sought to identify which of these methods remove batch effects and preserve biological variation best. Current approaches to detect batch effects involve visual inspection of dimension-reduced representations, such as principal component analysis (PCA). Yet, scRNA-seq data is high dimensional and sparse due to dropout events and stochastic gene expression, which may perturb the results of PCA^14^. In addition, it is unclear whether classical bulk transcriptome correction methods may need to be adapted to the sample-rich but sparse scRNA-seq situation.

Here, we propose a k-nearest neighbour batch effect test (*kBET*) to quantify batch effects in scRNA-seq data. Intuitively, a replicated experiment is well-mixed if a subset of neighbouring samples has the same distribution of batch labels as the full data set (**Fig. 1c**). In contrast, a repetition of the experiment with some bias is expected to yield a skewed distribution of batch labels across the data set (**Fig. 1d**). kBET uses a χ^2^-based test for random neighbourhoods of fixed size, followed by averaging the binary test results to return an overall rejection rate. This result is easy to interpret: low rejection rates imply well-mixed replicates.

In this study, we analysed five single-cell data sets derived from mice that cover both microwell plate-based and droplet-based methods with sample sizes ranging from 100 to 3000 cells per batch. We demonstrate the performance and accuracy of 11 normalisation and 5 batch effect regression approaches (**Fig. 1e**). Finally, we address the question of whether it is possible to integrate separate studies, and show with mouse development data sets that it is possible to correct for study-to-study effects. We show that batch correction based on *log(counts+1)*, *log(CPM+1)* or *scran* pooling, together with *ComBat* or *limma* regression, reduced the batch effect while preserving biological structure across all data sets (**Table 1**).

**Table 1:**
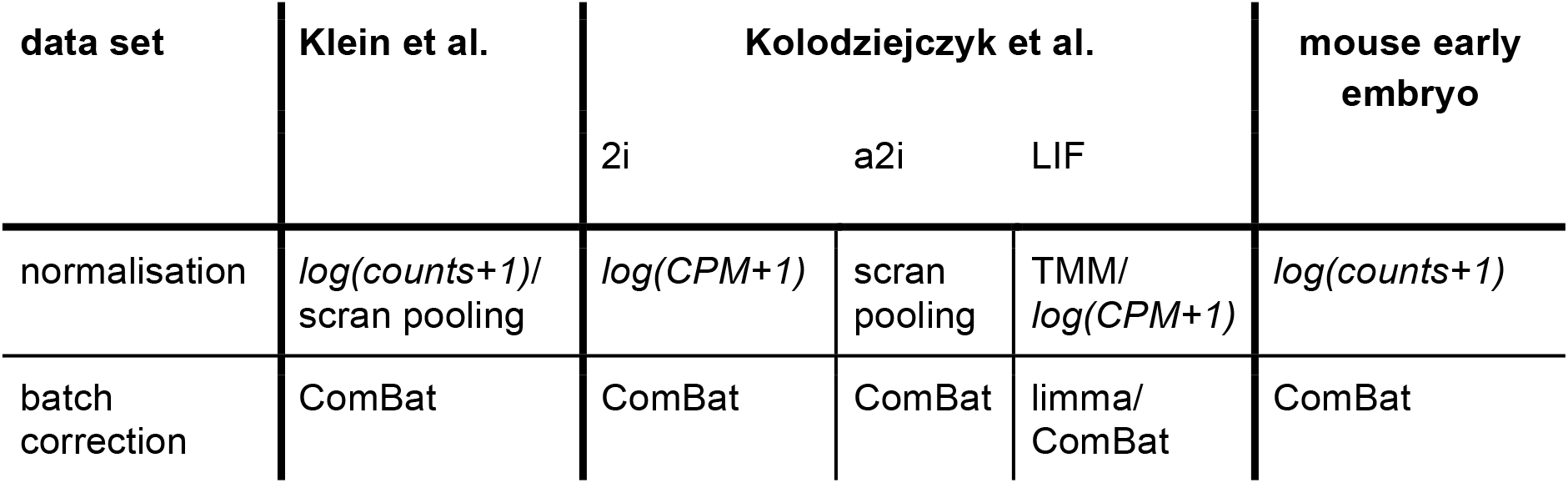
Best overall normalisation and batch correction methods. The ranking of batch correction strategies is based on kBET, retained HVG and false positive rates for Klein et al and Kolodziejczyk et al data. For mouse early embryonic development data integration, the ranking is based on both kBET and silhouette coefficient.

## Results

### kBET outperforms other methods for batch effect detection

We evaluated the performance of kBET on simulated data with three different degrees of batch effects: the data set consists of 500 samples (or “cells“) with 1000 “genes” each, where 1 %, 10 % or 20 % of the genes have their mean gene expression varied by a Gamma distributed random variable in the second batch (see **Methods** for details). Using appropriate scaling of this random variable, the expected mean gene expression remains unchanged. A batch with 1 % biased genes overlaps well with the other batch, yielding a low rejection rate (**Fig. 2a**). In contrast, a batch with 20 % biased genes separates from the other batch, so that samples/cells are surrounded by samples from the same batch only, yielding a high rejection rate (**Fig. 2b**).

**Fig. 2:**
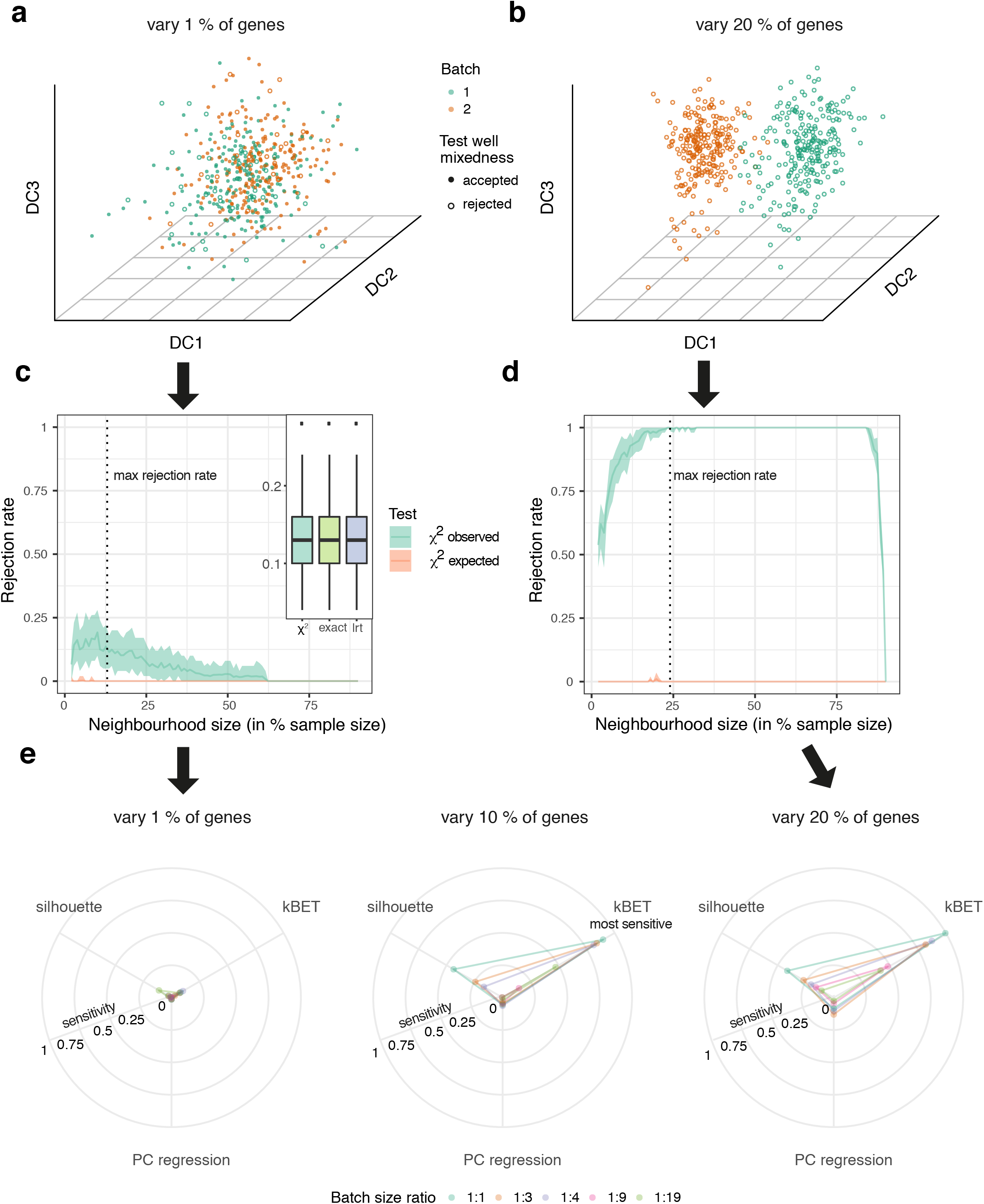
kBET is more sensitive than other batch tests on simulated data. Simulation results for 1000 genes and 500 cells. Two batches in a) and b) are equally sized with 1 % **(a)** and 20 % **(b)** varied gene mean expression levels, see **online Methods**. **c)** and **d)** illustrate how kBET values depend on neighbourhood size for a) and b). Dashed vertical line shows the optimal neighbourhood size for batch effect detection, *i.e.* where the rejection rate is maximal. Shaded areas represent the 95-percentile of repeated kBET tests (subset size 10 %). **e)** Comparison of kBET to other batch effect tests: PC regression and silhouette coefficient. Batch sizes were varied to assess the impact of unequal batch sizes.

kBET uses a Pearson’s *χ*^2^-based test for random neighbourhoods of fixed size *k* and averages the binary test result. Thus the neighbourhood size *k* is an important factor for the *χ*^2^-test that kBET is based on. For a small *k*, the rejection rate is smaller in general^15^. As soon as the neighbourhood size *k* for each test is larger than the size of a single batch, we observe a decrease in the rejection rate as well. This can be explained by the decreasing number of possible choices of batch labels; the ‘local’ batch label distribution becomes more similar to the global batch label distribution(**Figs. 2c** and **2d**). In between exceedingly small and large neighbourhood sizes, the average rejection rate becomes maximal. The value of the maximum indicates the presence of a batch effect (see **Supplementary Note 1**), and we use this maximum value for quantification.

kBET employs three different kinds of hypothesis tests: an exact multinomial test, Pearson’s *χ*^2^-test and a likelihood ratio test (*lrt*) (see **Methods** and **Supplementary Note 1** for details and ref. 15). The exact test yields an accurate result, but its computation is very costly as it involves the computation of each batch label configuration and the corresponding probability of observing it. Both Pearson’s *χ*^2^-test and a *lrt* approximate the result of the exact test with little deviation (inset in **Fig. 2c** and **extended Fig. 2**).

We compared the ability of kBET to detect batch effects with alternative measures: the average silhouette width (‘silhouette’) and principal component (PC) regression (**Fig. 2e**). In addition to the percentage of varied mean gene expression, we simulated different batch sizes ranging from equal size (1:1) to strong size imbalance (1:19). We found that kBET is most sensitive to the degree of bias compared with PC regression and silhouette. The silhouette performs better than PC regression, but silhouette shows little difference between 10 % and 20 % varied genes. kBET also performs better when only a few data points are biased by batch, as it still reveals a substantial bias when batches are imbalanced in size. Overall, kBET is clearly the most sensitive and robust measure of batch effect in this comparison.

### kBET accurately captures batch effects in single-cell RNA-seq data sets

Batch effects originate from different sources, as is evident when comparing technical replicates. We investigated the mouse embryonic stem cell (mESC) LIF cultures of Klein et al.^8^, which were generated with the inDrop protocol. The authors provided two technical replicates in the samples of day 0 culture (**Fig. 3a**), which offers an ideal case for batch correction assessment. The shift of the technical replicates in both PCs is a clearly visible inter-batch difference (**Fig. 3b**). We compared all combinations of normalisation and batch correction strategies, and illustrate f-scLVM corrected *log(CPM+1)* values (**Fig. 3c**) and *ComBat* corrected *log(counts+1)* (**Fig. 3d**). Both appear successful in removing batch effects according to visual inspection of the PCA. When we inspect each principal component, we find 196 PCs have a significant correlation with the batch covariate in the *f-scLVM* case, which explain 3.4 % variance in the data. For *ComBat* corrected *log(counts+1)*, none of the PCs correlates significantly with the batch covariate. The more sensitive kBET reveals that *ComBat* corrected *log(counts+1)* works best (**Fig. 3e**), in contrast to poor batch correction perfomance of *f-scLVM*. The PCA plot only shows the batch effect of the first two PCs, while kBET effectively quantifies more subtle batch effects.

**Fig. 3:**
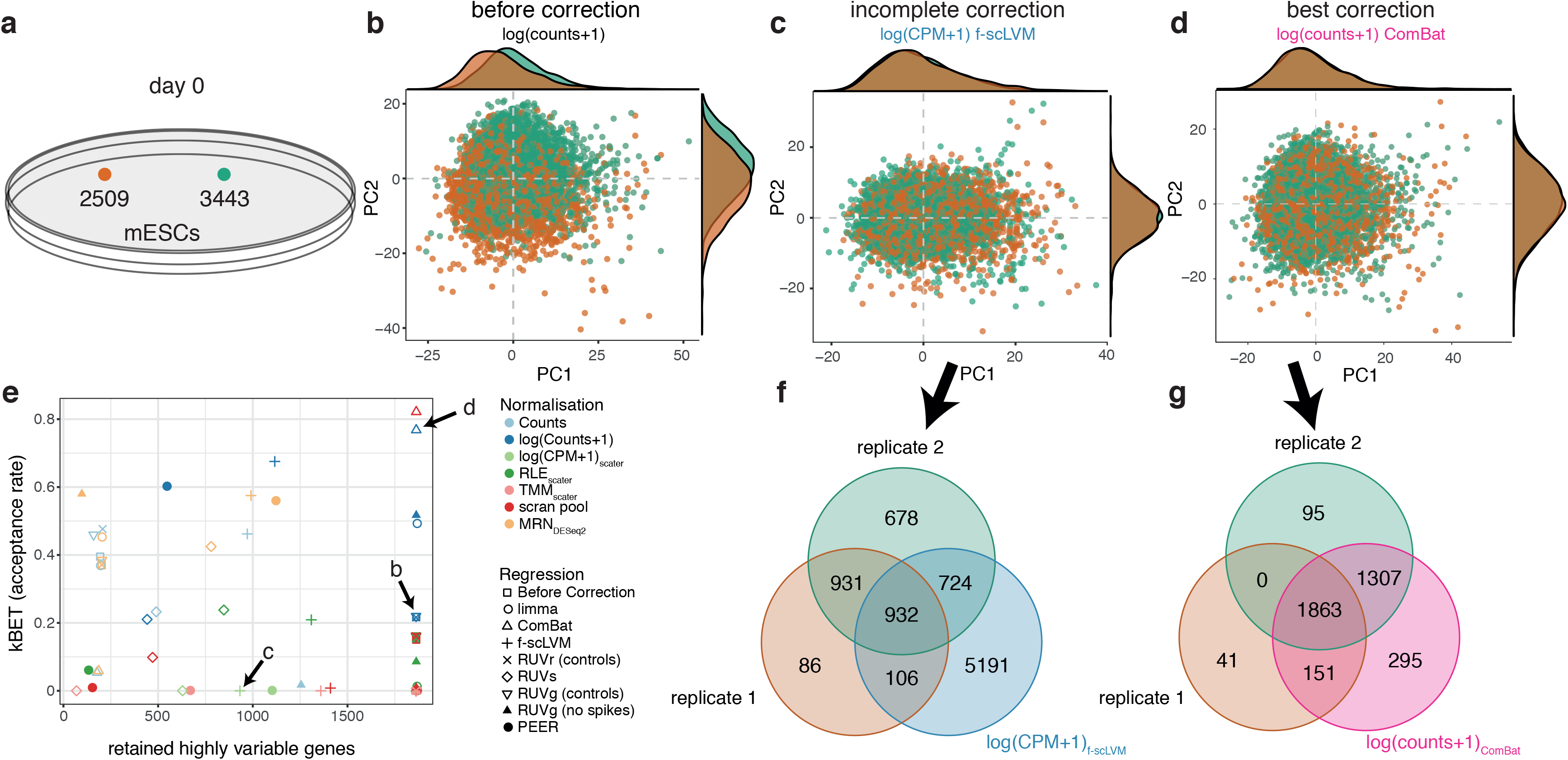
ComBat corrects best on mESC inDrop technical replicates (Klein et al.) The inDrop protocol provides a large UMI-count data set with two technical replicates **(a)**. PCA plots **(b-d)** display log-normalised counts, a biology-removing batch removal (f-scLVM on log-transformed CPM) and a biology-preserving batch removal (ComBat on log-transformed counts), respectively. Density plots depicted on the axes show the frequency of the replicates along the PCs. **e)** Percentage of retained highly variable genes vs. acceptance rate for all combinations of normalisations and batch regression approaches. **(f,g)** Venn diagrams of highly variable genes per replicate before correction and for the whole data set after batch correction. Highly variable genes in each replicate are computed on *log(counts+1)*values. The f-scLVM method retains 932 highly variable genes but has a high false positive rate, while ComBat captures all the highly variable genes with a low false positive rate.

### Distinguishing batch effect variability from biological variability

The second challenge in batch correction is to preserve biological heterogeneity in the data, otherwise the optimal batch correction would remove all variance, setting each observation to the same constant. We assess biologically relevant heterogeneity by computing highly variable genes (HVG) before and after correction. Before correction, we only consider HVG present in all replicates. (We find considerably more HVG in the whole data set than replicate-wise, due to the batch effect.) For example, let A, B and C represent three replicates, and *a*,*b* and *c* the corresponding sets of HVG. Then, the batch-free, conservative set of HVG is the intersection of *a*,*b* and *c*: *HVG*_*batch−free*_ = *a* ∩ *b* ∩ *c*. When we check the set of HVG after batch correction (HVG_corr_), this set would ideally contain the complete set of HVG_batch-free_. In total, we evaluated the fraction of retained HVG after correction (see **Methods** and **Fig. 3f-g**).

To complement the concept of retained HVG, batch correction is not supposed to introduce additional variability to the data. Thus, we consider the difference set of HVG before and after correction, i.e. *HVG*_*corr*_\(*a* ∪ *b* ∪ *c*) as false discoveries that we use to compute a false positive rate (FPR, see **Methods**). Here, the two technical replicates share 1863 HVG_batch_free_ and over 700 HVG reside in either of the replicates (**Fig. 3f-g**).

After correction by *f-scLVM*, we retained half of HVG_batch-free_ while we discovered over 5000 HVG in the whole data set (**Fig. 3f** and **extended Fig. 3a-b**), which explains *f-scLVM*’s minimal kBET acceptance rate (**Fig. 3e**). When we compute the FPR on the basis of *log(CPM+1)* normalised data, we find a FPR of 27 % (**extended Fig. 3c**). We obtained the best result for the combination of log-transformed Counts and *ComBat* (**Fig. 3d**) - all HVG_batch-free_ were kept after batch correction and only 295 HVG were caused by batch correction (8 % FPR, see **Fig. 3g**).

In conclusion, batch correction may confound observations massively, masking the biological signal completely. In addition, even the best batch correction strategy leaves part of the batch effect in the data (**Fig. 3e, g**). This explains the increase of the total amount of HVG after correction (**extended Fig. 3b**) and FPR (**extended Fig. 3c**). Both the silhouette coefficient and PC regression show little discrimination between most of the correction strategies (**extended Fig. 3d-e**), whereas kBET resolves them in detail (**Fig. 3e** and **extended Fig. 3d-e**).

### Best practice in batch correction

Next, we examined mouse embryonic stem cells cultured in three different conditions (2i, a2i and LIF)^16^ and sequenced with the SMARTseq2/C1 protocol (**Fig. 4a**). These data sets are rather similar in terms of heterogeneity, but the biological origin of the heterogeneity in each culture condition is different (compare ref. 16 for details). Thus applying the same batch correction and normalisation strategies led to similar results across all culture conditions. We obtained well-mixed data for all data sets with *log(CPM+1)* normalisation and batch correction with *ComBat* (**Fig. 4b** and black arrows in **Fig. 4c**).

**Fig. 4:**
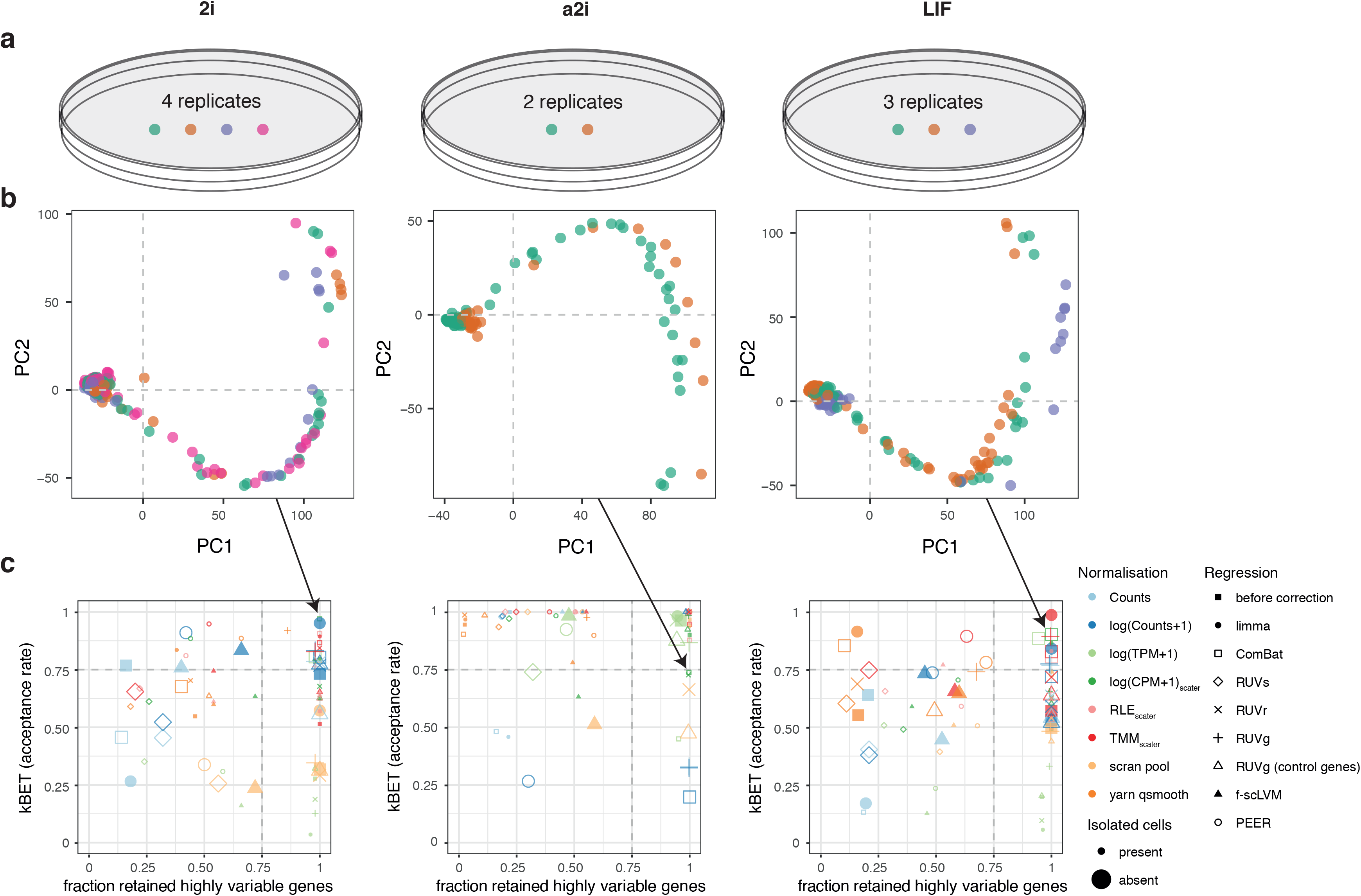
Deeply sequenced SMARTseq2/C1 mESC data have similar characteristics for batch correction (Kolodziejczyk et al.) **(a)** Illustration of three full-length read data sets with replicates in 2i, a2i and LIF culture (219, 123 and 207 cells, respectively). **(b)** PCA plots for *log(CPM+1)* ComBat corrected data. **(c)** Percentage of retained highly variable genes vs. kBET acceptance rate for all combinations of normalisation and batch correction approaches. Best performing normalisation-regression strategies cluster in the top right corner, such as ComBat on *log(CPM+1)* data.

In all data sets, log-transformed count data are locally well-mixed in over 50 % of cases (dark blue full squares in **Fig. 4c** and **extended Fig. 4a-c**). We notice a decrease in acceptance rate in 2i and LIF culture when we compare count data to other normalisation methods. For batch effect correction, *RUV* controlled both acceptance rate and overlap in highly variable genes, but we also noticed an increase in the total number of HVG (**extended Fig. 4d-f**) as well as an increased false positive rate (**extended Fig. 4g-i**). In contrast, *ComBat* increased the acceptance rate, preserved almost all HVG_batch-free_ in all data sets and had a low false positive rates at the same time.

In a2i, we observe almost perfect mixing in log-transformed data, but we also found that a considerable amount of cells are *isolated*, *i.e.* they do not have a mutual nearest neighbour. These cells are implicitly removed from the local structure evaluation. If the isolated cells have a different label composition than the global data set, we find strong differences in kBET’s rejection rate if isolated cells remain unconsidered (see **extended Fig. 4j-k**). Therefore, we adapt the expected label composition such that we can neglect the isolated cells (see **Methods**). In general, we consider a correction strategy is ill-advised if it produces considerable amounts of isolated cells when removing batch effects (see **extended Fig. 4a-c**).

The performance of batch correction methods varied slightly from data set to data set (**Fig. 4c**). To test the dependence of batch correction performance and the number of batches, we subset the 2i data to combinations of three and two batches (see **Supplementary Note 2** and **Supplementary Fig. 4b,c**). In these batches, library size and number of detected genes do not correlate well (**Supplementary Fig. 4d**). Depending on the combination of batches, we observed performance differences in batch effect correction that was independent on the number of batches and rather explicable by batch-specific dissimilarity. Taken together, we demonstrated that classical batch correction tools, in particular *ComBat*, successfully remove batch effects and preserve the biological signal (**Table 1**).

### Going beyond replicates - data set integration across multiple studies

With the explosion of single cell RNA-seq data in recent years^17^, we begin to realise the need for a comprehensive strategy of data integration. Of course, correcting batch effects between different studies is more challenging than within the same study, especially, if cell types vary between studies. In this work, we benchmark the batch correction performance on 8 different Smart-seq based data sets^18–25^, all profiling mouse early embryonic development from oocyte to blastocyst (**Fig. 5a** and **online Methods**).

**Fig. 5:**
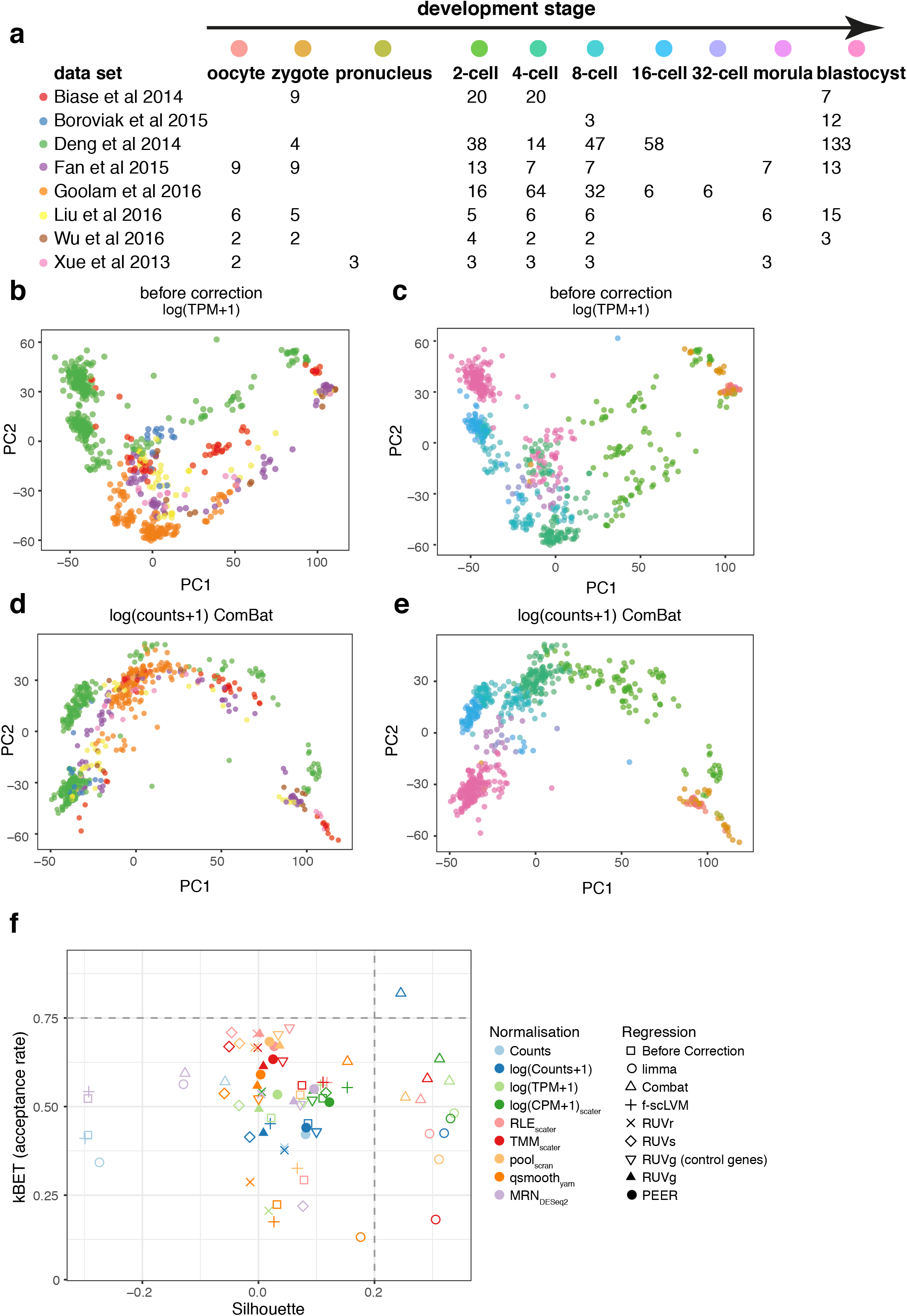
kBET assesses in agreement of cell stages in early mouse embryo data integration. **(a)** Overview of data from early mouse development from 5 different data sets. **(b-c)** PCA plots of *log(counts+1)* normalised expression data coloured by data set (b) and by developmental stage (c). **(d-e)** PCA plots of of *log(counts+1)* normalised expression data after batch correction by limma coloured by data set (d) and by developmental stage (e). **(f)** Silhouette coefficient of embryonic development vs. average kBET acceptance rate (weighted per developmental stage) reveals that *ComBat* applied to *log(counts+1)* provides good mixing of cells from different studies in the same developmental stages. This is indicated by the high kBET acceptance rate and by high silhouette coefficient, indicating the best separation of developmental stages.

We remapped the reads to the same reference transcriptome with Salmon^26^ to reduce quantification biases^27^. Interestingly, even different versions of Salmon resulted in different degrees of batch effect (see **Supplementary Note 3**). Batch effects before correction are quite obvious even in PCA (**Fig. 5b-c**): Biase et al data and Deng et al data deviate significantly from others. Consequently, the cells are more likely to cluster by study rather than embryonic developmental stage. However, we observe that it is possible to correct the batch effect computationally: the best results are with *ComBat* on *log(counts+1)* (**Fig. 5d-e**) with an average acceptance rate of 82 % (**Supplementary Table 2**).

When integrating developmental data, we expect that the same cell types from different studies mix, maintaining the correct trajectory of successive developmental stages. Hence, we assess the batch effect of each developmental stage based on averaged kBET results. In parallel, we monitor the developmental progression by silhouette (**Fig. 5f)**. A high acceptance rate implies good mixing within each developmental stage, and a good separation of developmental stages translates into higher silhouette coefficients. Before correction the developmental stages separate weakly (silhouette 0.08 for *log(counts+1)*). After correction with Empirical Bayesian methods as *limma* and *ComBat*, we observe distinct clustering according to development stages, but only *ComBat* achieves a good mixing of study batches. Notably, the PC1 corresponds to the real developmental time of the cells.

As desired, one of the normalisation and regression methods captures both the developmental stages and mixes the data from different studies. *ComBat* with *log(counts+1)* yields high acceptance rates and clear clustering by developmental time. This example illustrates how batch effect correction tools can play a key role in data integration and provide an effective separation of the biological signal from complex technical variations. In this work, we considered each study as one batch and ignored any technical substructure within the studies. Also, cell type distribution as well as cell collection time points differed slightly across studies, which made the task more difficult. For future data integration efforts with more complex data structure and less prior knowledge on cell types, the community needs more sophisticated batch correction methods that model nested batch structures and several batch variables.

## Discussion

Batch effects in single-cell RNA-seq data can have a severe impact on downstream data analysis if they are not properly accounted for. Moreover, they have a substantial random noise component that stems mostly from technical factors of the experiment. In the simplest possible case, where we have technical replicates that are otherwise homogeneous, *ComBat* corrects the data and preserves the underlying biological properties (**Supplementary Table 2**). At the next level of complexity, with biological replicates such as two independent cell cultures of the same cell type and more batch-to-batch variability, *ComBat* again dealt well with the situation.

Current batch correction methods have been designed to correct bulk RNA-seq or microarray data. With little or no adaptation, they can be applied to single-cell RNA-seq data. While single-cell RNA-seq data mirror cell-to-cell variability, they are sparse because of dropouts in the experiment. Yet, none of the current batch effect correction approaches tackles the dropout property in single-cell RNA-seq. (A mere mean shift and variance stabilization would not take into account a batch-to-batch difference that is solely addressing dropout rates.) Moreover, with thousands of measured cells per data set, optimal memory usage and efficient implementation will be as important as accurate correction for confounders (**Supplementary Note 4**).

In contrast to batch correction with regression models, normalisation aims to reduce cell-to-cell bias within a batch. Previous studies have discussed appropriate scaling factors ^2,6^, but we found that normalising for library size with CPM consistently increased batch effect compared to raw count data. Also, the number of genes per cell and the library size may not correlate well across batches (**Supplementary Fig. 4d**). Nevertheless, CPM normalisation and the more advanced scaling with scran in combination with *ComBat* worked very well in deeply sequenced SMARTseq2/C1 data (**Fig. 4** and **Table 1**). In addition, all batch effect correction methods require a certain statistical property in their model. For example, *RUV* requires count data with negative binomial distribution. If the data input in *RUV* violates the model assumption, *RUV* introduces additional variability to the data, which we described as isolated cells.

The k-nearest neighbour batch effect test (kBET) approach allows the study of high-dimensional data without prior assumptions regarding statistical properties. Hence, kBET is applicable to any type of NGS data given a reasonable sample size per batch. Still, the underlying model assumption requires all batches to be equivalent and interchangeable. While simple, the translation into balanced experimental design is challenging. For complex experimental setups as time series data collection, it would require one to collect and sequence all cells of all time points together. Otherwise data is confounded with both technical factors and biological variation between samples. With the imminent global efforts to create a genomic single-cell reference of every tissue in the human body - the Human Cell Atlas - we need to be able to robustly determine, control and correct numerous sources of both technical and biological variations.

## Methods

Methods, including statements of data availability and any associated accession codes and references, are available in the online version of the paper.

## Acknowledgements

We would like to thank Anika Böttcher for motivating this study with data, and Tomislav Illicic for carrying out pilot analyses. We are grateful to the members of the Teichmann and Theis labs for valuable discussions and comments on the manuscript.

## Online Methods

### kBET -- *k*-nearest neighbour batch estimation

Let the full gene expression data set *D* = {*x*_1_, …, *x*_*n*_}, where *x*_1_ ∈ ℝx^*g*^ and and *X* ∈ ℝ^*n×g*^ the corresponding gene expression data matrix with *n* samples and *g*genes. In a single-cell RNA-seq data set *X*, each sample has meta-information such as cell type, FACS gate or the batch *i* it was measured in.

The batch variable *i* has *l* categories such that *n*_*i*_ denotes the number of samples in batch *i*, 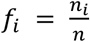 the fraction of samples in *i*, and *v* = (*n*_1_, …, *n*_*l*_) the *batch configuration* of all samples.

We formulate the null hypothesis of “data being *well-mixed*'”, *i.e.* the absence of a batch effect, as

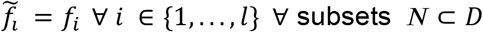

In order to statistically test this hypothesis, let us define a *neighbourhood* subset *N*_*j*_ = x_j_ ∪ {*x*_*s*_ ∣ *s* is among *k* − 1 nearest neighbours of *j*}. Nearest neighbours are computed with the *cover-tree* algorithm (*FNN* R package). To optimize computation efficiency, we pre-compute the first 50 eigenvectors of the largest eigenvalues with the *svd* function and use the reduced data set to find nearest neighbours.

Let *n*^*k*^_*ji*_ denote the number of cells in batch *i* that are in subset *j* of size *k*. Testing the null hypothesis involves two steps:

1. We test the null hypothesis in each subset *N*_*j*_ of a given sequence of subsets. In each subset *N*_*j*_, this amounts to testing whether the distribution of *n*^*k*^_*ji*_ with respect to *i* matches the distribution under the null hypothesis.
2. We summarize the result of the sequence of tests by computing the average rejection rate *S* over all tests -- a test statistic for the whole data set. Hence, testing whether *S* exceeds a significance threshold allows to reject the null-hypothesis for the whole data set.

Note that by performing these two steps, we go beyond a standard test for *homogeneity* of subsets of a given data set.

#### *χ*^2^-based test

In the limit of high values of *k*, *n*^*k*^_*ji*_ is Gaussian distributed with respect to *i*. A test for small values of *k* is provided as exact test (**Supplementary Note 1**). Then, we can use Pearson's *χ*^2^ test, the test statistic of which reads

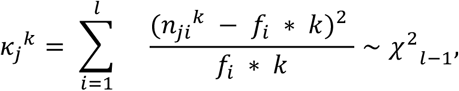

where *χ*^2^_*l-1*_ denotes the χ^2^-distribution with *l-1* degrees of freedom. The p-value for each *k*_*j*_^*k*^ is computed as

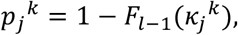

where *F*_*l−1*_(*x*) denotes the cumulative distribution function of the χ^2^-distribution with *l-1* degrees of freedom.

#### Identification of isolated cells and adaptive frequencies

When we determine the local structure of the data as neighbourhoods, we implicitly assume that every cell has mutual nearest neighbours. That means we expect that every cell can be found in more than one neighbourhood of size *k*. If a cell is more distant to its *k-1* nearest neighbours than those neighbours among each other, we call this cell an *isolated cell* or *outsider*. Such a cell does not contribute to any other neighbourhood and the composition of a tested neighbourhood will not change if an *isolated cell* is removed from the data entirely. However, a considerable amount of isolated cells bias the observed outcome if their label composition is significantly different from the global batch label distribution. We introduce adaptive expected frequencies that are computed without the isolated cells to adjust for this effect.

Let *N*_*iso*_ = {*x*_*j*_∣ *j*has no mutual nearest neighbour} and *n*_*i*_^*iso*^ denotes the number of cells in batch *i* that are isolated. Then, we apply Pearson’s χ^2^ test to determine if a certain batch label class is overrepresented:

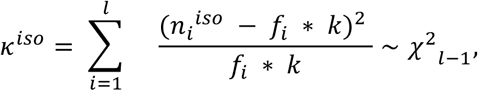

where the p-value reads *p*^*iso*^ = 1 − *F*_*l-1*_(*k*^*iso*^), where *F*_*l-1*_(*x*) denotes the cumulative distribution function of the χ^2^-distribution with *l-1* degrees of freedom. In case that *p*^*iso*^ < α = 0.05, we compute adapted expected frequencies on the basis of *v* = (*n*_1_ − *n*_1_^*iso*^ … *n*_l_ − *n*_l_^*iso*^) the *batch configuration excluding isolated cells*.

### Principal component regression

Principal component analysis (PCA) is an orthogonal transformation the data matrix *x* ∈ *R*^*n×*^ into a set of linearly uncorrelated variables. The principal components (PCs) correspond to the eigenvectors of the covariance matrix *Cov*(*X*) of the data and are ordered by the explained variance of the data. If a batch effect is present in the data, it contributes to the variance. As the set of PCs is uncorrelated, regressing the batch covariate *B* (with *l* categories defined in the kBET model and the *i*^*th*^ PC returns the coefficient of determination *R*^2^(*PC*_*i*_|*B*)as approximation of the variance explained by *B* in each PC (principal component regression, similar to ref. 28). Overall, the total contribution of the batch effect to the variance in the data may be approximated by

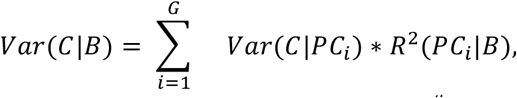

where *Var*(*C*|*PC*_*i*_) is the variance of *C* explained by the *i*^*th*^ PC. However, using a linear regression model enables us to evaluate the significance of *R*^2^(*PC*_*i*_|*B*).For the case of two batches, the significance test equals a univariate t-test on the loadings of each PC split by batch covariate. However, as the number of features (genes) increases, the largest and smallest eigenvalue of the sample covariance matrix converge ^29^. Consequently, *Var*(*C*∣*B*) decreases with the number of features as well and due to the high-dimensionality of scRNA-seq data, batch effects are underestimated.

We use the sum of explained variance of the PC with the most significant (i.e. highest) *R*^2^(*PC*_*i*_∣*B*)as proxy for the batch effect:

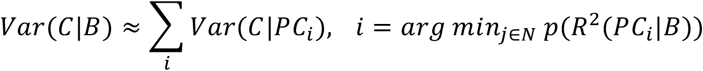

### Silhouette coefficient

The calculation of a silhouette aims to determine if a particular clustering has minimised within-cluster dissimilarity and maximised inter-cluster dissimilarity^30^. Let us assume there is a given clustering into more than one cluster. For each sample *i*, the silhouette width *s*(*i*) is defined as follows:

Let *a*(*i*) be the average dissimilarity between *i* and all other data points of its cluster A. If *i* is the only observation in this cluster, set *s*(*i*): = 0. For all other clusters *C* ≠ *A*, let *d*(*i, C*) be the average dissimilarity of *i* to all samples of C. There is some cluster B whose dissimilarity *d*(*i,B*) is minimal: *b*(*i*): = *min*_*c*_*d*(*i,C*), which is the "neighbouring" cluster to sample *i*. Then, the silhouette width *s*(*i*) is defined as the scaled difference of average dissimilarity within a cluster and the average dissimilarity to its "neighbouring" cluster:

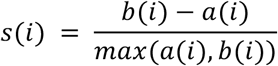

Finally, the mean of all silhouette widths *s*(*i*) gives the silhouette coefficient *s* from which we display its absolute value (in **Fig. 2**). We adapted the calculation from the *scone* R package^5^.

### Computation of highly variable genes

In order to determine if a batch correction method is over-correcting, we check the number highly variable genes (HVG) before and after batch correction. In the Brennecke^31^ model implemented in the M3Drop^32^ package, the relation of the squared coefficient of variation (CV^2^) and mean μ for each gene follows a Gamma model 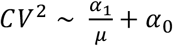 The CV^2^ decreases with increasing gene mean expression. A gene is considered as highly variable, if its CV^2^ is higher than expected from its mean.

To define a batch-free gene set before batch correction, we fit the Brennecke model to each batch separately and intersect the corresponding sets of HVG. Let *l* be the number of batches and a_i_ the set of HVG for batch *i*, then we denote

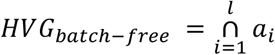

as the set of HVG present in each of the batches in a data set.

More specifically, we considered the fact that highly variable genes depend on the type of normalisation^6^. Then, the reference set of highly variable genes consists of all genes that are highly variable in all batches with *log(counts+1)* normalisation. After batch correction, we compute HVG for the whole corrected data set (HVG_corr_). Ideally, we would retain all HVG_batch-free_ after batch correction. We define the fraction of retained batch-free HVG, 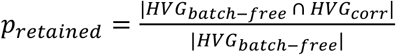 to determine if the biological signal in the data is preserved upon batch correction.

### False positive rate for highly variable genes

We quantify the number of HVG caused by the batch effect as a false positive rate (FPR). In contrast to the fraction of retained HVG, we define the FPR by the fraction of HVG that are found in the whole data set but not in any of the batches. More formally, let

- a: set of highly variable genes in the complete data set and
- a_i_: set of highly variable genes in batch *i*.

Then, the false positive rate reads

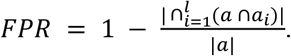

### Data Normalisation

Data normalisation methods account for the sequencing depth as a size factor and normalise the expression data to the same comparable level. We summarised the normalisation methods used in **Supplementary Table 1.** Briefly, 1) Counts per million (CPM) is based on the library size; 2) Relative log expression (RLE); 3) Trimmed Mean of M-values (TMM); 4) scran size factor^13^; 5) qsmooth from the *YARN* package^33^; 6) Transcripts per million (TPM) is derived from the mapping by Salmon^26^(version 0.8.2).

### Batch regression

Methodologically, the recent batch regression approaches either require the assignment of batches as input or they assess bias in the data independently from batch information. In this paper, we compare five established batch regression methods (see **Supplementary Table 1** for details).

1) *limma* employs an Empirical Bayes model, we used the *removeBatcheffect* function from the *limma* package^34^; 2) the Combat model^35^ function from *sva* package^36^, which is based on Empirical Bayes methods; 3) the f-scLVM model is a factor analysis based latent variable model, after training the model, the batch effect related factors or removed using the regressOut function implemented in the *fscLVM* package^37^; 4) PEER is based on factor analysis^38^; 5) RUVs, RUVr and RUVg from *RUVseq* package^5^ remove unwanted variance according to replicate samples, residuals and control genes. We derive control genes using the *edgeR* package^39^ and the top 400 constant genes are used as control genes. The model parameter *k* in RUVseq and PEER indicates the number of hidden factors correlated to the variance. We tested several values from 1 to 7 and 25 % of the sample size. Methods 1-3 require batch information for correction, methods 4-5 assess general bias in the data.

### Simulated data

We model the number of transcripts per gene and per cell as count data that follow the negative binomial distribution with zero-inflation (ZINB) to account for dispersion and sparsity caused by dropouts. Mean expression levels for each gene are sampled from the beta-distribution (with appropriate scaling):

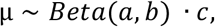

with parameters *a* = *2*,*b* = 5 and *c* = 100. The dropout probability for each simulated gene *j* ∈ {1,…, *G*} in batch *i* ∈ {1,2} is modeled by the logistic (sigmoid) function *p*_*ij*_ = *sigm*(−(*β*_0_ + *β*_1,*i*_*μ*_*ij*_)), where *β*_0_ = −1.5 and *β*_1,*i*_ = 1/*median*(*μ*_*i*_). In total, every sample is drawn from *s*_*ij*_ ∼ *NB*(*μ*_*ij*_, *θ* | *Ber*(*p*_*ij*_)), where *θ* = 1 and *Ber* is the Bernoulli distribution. The mean expression levels of the second batch *μ*_2_ are subject to different degrees of variation. We multiply a 1 %, 10 % and 20 % of the mean expression levels *μ* with a Gamma distributed random variable *γ* ∼ *Г* (*α, β*) and *α* = *β* = 1:

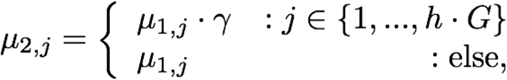

where *h* ∈ {1 %, 10 %, 20 %} and *G* is the number of genes in the data set. The Gamma distribution with the chosen parameters has mean and variance equal to 1 such that the expected value of the sampled mean expression levels stays unchanged. In addition, we vary the sample size of the two batches: In each simulation, we sample 500 instances with 1000 genes each, with the size ratio of the batches being 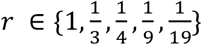 This means equally sized batches contain 250 samples each, and batches with 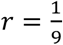 have 450 and 50 samples, respectively.

### Public data sets

We applied the batch estimates to several single-cell RNA-seq data sets. In the inDrop publication, the droplet based sequencing was demonstrated on mouse embryonic stem cells growing on LIF^+^ medium and additional two technical replicates^8^. In our analysis, we have used the two replicates that consist of 5952 cells from two batches and 11308 genes with at least 2 cells having more than 4 UMI reads per cell.

Kolodziejczyk et al.^16^, explored heterogeneity in mESCs cultured with three different media (2i, alternative 2i and LIF^+^) on full-length sequenced transcripts (SMART-Seq). The three conditions include 219, 123 and 207 cells on 4, 2 and 3 batches, respectively.

Further, single-cell RNA-seq has been widely applied in exploring mouse embryonic development. To test the performance of batch correction for data integration, we collected single cell RNA-seq data of mouse early embryonic development from 8 different studies^18–25^, which consist of 56, 49, 124, 65, 15, 294, 17 and 15 cells, respectively. The corresponding numbers of cells per cell developmental stage are summarised in **Fig. 5a**.

### Data sources

The mESC data sequenced with inDrop^8^ were downloaded as UMI-filtered read count matrices with accession number GSE65525.

The mESC data sequenced with full length SMART-seq^16^ were downloaded from ENA (project id: PRJEB6455) as fastq files and mapped to Ensembl^40^ mouse transcriptome (GRCm38.p5.87, equivalent to UCSC mm10) with Salmon^26^. Cells were quality controlled according to data derived from the Espresso database (<u>http://www.ebi.ac.uk/teichmannsrv/espresso/</u>).

Early embryonic development data were derived from several studies^18–25^ with accession ids: E-GEOD-57249, E-GEOD-70605, E-MTAB-3321, GSE53386, E-MTAB-2958, E-GEOD-45719, E-GEOD-44183 and E-GEOD-66582. All studies applied SMARTseq-based protocols for single-cell RNA-seq. All fastq files were mapped to Ensembl^40^ mouse transcriptome (version GRCm38.p5.87) with Salmon^26^ (version 0.8.2, kmer = 21 to tolerate different read length). Here, we only consider the studies as batches while omitting the flowcell batches. We continued our analysis without further gene filtering or quality control.

### Software availability

kBET is available as an R package at <u>https://github.com/theislab/kBET</u>.

An implementation of the batch regression methods is available at: <u>https://github.com/chichaumiau/batch_regression/</u>.

